# Mapping groundwater-dependent vegetation in temperate climates on the example of Central Germany

**DOI:** 10.64898/2026.03.12.706590

**Authors:** Léonard El-Hokayem, David Emanuel Schulz, Christopher Conrad

**Affiliations:** Institute of Geosciences and Geography, Martin Luther University Halle Wittenberg, Halle (Saale), 06120, Germany; German Centre for Integrative Biodiversity Research (iDiv) Halle Jena Leipzig, Leipzig, 04103, Germany

**Keywords:** Groundwater-Dependent Vegetation, Remote Sensing, Biodiversity, Phreatophytes, Machine Learning, EU Waterframework Directive

## Abstract

Groundwater-dependent ecosystems are biodiversity hotspots that provide habitat for specialised species. The EU Water Framework Directive (WFD) stresses the importance of identifying and protecting these ecosystems. However, they remain poorly mapped in temperate regions, as most studies have focused on (semi-) arid regions, where groundwater use by vegetation is both more prevalent and easier to detect from remote sensing. In this study, we transfer mapping approaches for groundwater-dependent vegetation (GDV) from dry climates into a novel framework for humid climates. To do so, we integrated, ECOSTRESS evapotranspiration data, together with high-resolution remote sensing data, regional geospatial data and field data to identify GDV. To test our framework, we trained and validated Random Forest models with eight predictor variables using 166 ground-truth vegetation plots to map GDV in Saxony-Anhalt (Germany). The final model achieved an overall accuracy of 0.97, identifying 2,067 km^2^ (41%) of GDV. Currently, only 19% are protected under the EU WFD. The proposed mapping framework offers a new solution for identifying GDV in temperate regions. The new GDV maps can contribute to managing groundwater resources and preserving biodiversity hotspots in regions facing increasing droughts, ultimately supporting implementation of the EU WFD.

## 1 Introduction

Protecting groundwater-dependent ecosystems in accordance with the EU Water Framework Directive (WFD) (EC, 2000) requires knowing where these ecosystems occur. As droughts intensify across Europe (Gebrechorkos et al., 2025), mapping these ecological refugia is increasingly urgent. Still, many groundwater-dependent ecosystems remain unmapped globally, particularly outside dryland regions (El-Hokayem et al., 2026; Rohde et al., 2024b).

Groundwater-dependent vegetation (GDV) comprises plant communities that rely on groundwater to sustain their composition, structure, and function (Eamus et al., 2015; Barron et al., 2014). These include both groundwater-fed surface ecosystems, such as spring and wetland vegetation, and phreatophyte communities accessing subsurface groundwater (Eamus et al, 2015). Phreatophytes are plant species capable of accessing groundwater that often act as ‘foundation species’ and indicators of GDV presence (Rohde et al., 2024a; Canadell et al., 1996; Meinzer, 1927). In temperate regions, GDV is typically associated with riparian zones, wetlands, and groundwater-fed forests (Bug et al., 2021; Bhatnagar et al., 2020), where groundwater availability is controlled by local climate, geology, and topography (Doody et al., 2017).

GDV supports regional biodiversity, provides refugia and stepping-stone habitats, and contributes to ecosystem services such as flood mitigation, water purification and groundwater recharge (Brown et al., 2009; Kløve et al., 2011). However, declining groundwater levels due to climate change, land-use intensification, and overexploitation increasingly threaten the resilience of these systems globally (Eamus et al., 2016; Kløve et al., 2014). Policy frameworks, such as the EU WFD address this vulnerability, and highlight the necessity to map and protect GDV (Baaner et al., 2025; EC, 2000). Yet, GDV have rarely been systematically mapped at larger scales relevant for conservation or management across temperate Europe (Kilroy et al., 2008). This gap underscores the need for robust spatial tools that can support both biodiversity protection and EU WFD implementation.

Despite their global importance and occurrence (Link et al., 2023), GDV research remains geographically biased. Most studies focused on arid and semi-arid biomes, where groundwater dependence is both more critical for vegetation productivity and stability and easier to detect (Glanville et al., 2023; Rampheri et al., 2023). In temperate regions, studies on GDV identification remain scarce, even though groundwater subsidises tree growth and transpiration (Ciruzzi & Loheide II, 2021; Peters et al., 2008) and the overall uptake by vegetation is estimated to be 18%–30% (Link et al., 2023; Evaristo & McDonell, 2017). This spatial bias limits ecohydrological insight and hinders the development of spatially explicit management for temperate GDV.

Precise identification of GDV mainly relies on field-based methods, such as vegetation surveys, isotope analysis or hydrogeological sampling. However, these methods are generally resource-intensive and difficult to scale (Koit et al., 2021; Kilroy et al., 2008; Batelaan et al., 2003). At the regional level, geospatial data, such as biotope types or groundwater table depth, is often overlaid to map at least GDV potential (Duran-Llacer, 2022; Doody et al., 2017). Although remote sensing offers more precise and scalable approaches to mapping GDV, these have mainly been developed and tested in dry regions. (Pérez Hoyos et al., 2016), where the identification of persistent ‘green islands’ during seasonal droughts is widely applied (Akasheh et al., 2008). Several studies suggest that such frameworks are transferable to other climates (Rohde et al., 2024b; Gou et al., 2015), but this transferability has never been tested. However, the absence of annual dry seasons in temperate regions makes it difficult to detect drought-related responses in vegetation (Brim Box et al., 2022). Consequently, mapping GDV in temperate regions requires predictor variables and models tailored to temperate climatic conditions and vegetation types. Evapotranspiration (ET) is a key indicator for identifying GDV as it directly reflects plant water use, drought response, and ecosystem health (Fisher et al., 2020). Prior studies used MODIS or Landsat imagery together with hydrological models to estimate ET of GDV (Gow et al., 2016; Adams et al., 2015; Yang et al., 2011). The ECOSTRESS mission offers ET data with higher temporal resolution, which has been successfully applied across ecosystems (Kohli et al., 2020; Liang et al., 2022; Xiao et al., 2021). However, despite its potential, ECOSTRESS ET data has yet to be applied in GDV mapping, representing a novel opportunity to enhance detection, quantify water use and improve mapping frameworks.

In this study, we present the first high-resolution remote sensing-based framework for mapping GDV in temperate regions. Building on approaches developed for semi-arid regions (e.g., El-Hokayem et al., 2023a; Gou et al., 2015), we adapt and extend them by integrating for the first time ECOSTRESS ET data together with Sentinel-2 vegetation indices, hydrogeological data, and topographic predictors. Botanical field surveys of 171 vegetation plots in Saxony-Anhalt (Germany) provided ground-truth data to train and validate regional Random Forest models. Existing biotope maps allowed for plausibility analysis of the outcomes. Finally, we test the framework to produce the first high-resolution GDV map for a study region in Central Germany.

Our study follows three objectives: I) To adapt and test GDV mapping frameworks developed in semi-arid and arid regions for use in temperate regions. II) To integrate satellite-derived ET data from ECOSTRESS with multispectral vegetation indices, hydrogeological, and topographic predictors and ground-truth vegetation data for GDV mapping. III) To produce and validate the first high-resolution GDV map for the study region of Saxony-Anhalt, Germany. By addressing these objectives, this study delivers both a novel framework for temperate GDV detection and a regional application that supports science-policy integration in groundwater and ecosystem management within the EU WFD.

## 2 Material and methods

The developed workflow for mapping temperate GDV consists of three steps: i) adapting existing frameworks originally designed for semi-arid areas (El-Hokayem et al., 2023a; Gou et al., 2015), ii) applying the adapted framework to the study area of Saxony-Anhalt (Germany), and iii) validating outcomes using field survey data and biotope maps. To do so, the framework combined ground-truth vegetation plot data with predictor variables derived from remote sensing and geospatial datasets (Supplementary 1 for workflow and details).

### 2.1 Study area

Situated within the ‘Temperate Broadleaf & Mixed Forests’ biome (Dinerstein et al., 2017), the study area encompasses the federal state of Saxony-Anhalt, Germany. The region has a warm-summer humid continental climate (Dfb), with a mean annual temperature of 8.7 °C and an average annual precipitation of 548 mm (1991–2021) (Kronenberg et al., 2021). Saxony-Anhalt lays in the Central German Dry Region, where annual precipitation can fall below 450 mm, due to the rainshadow of the Harz Mountains, making it the driest German state. Combined with high ET, this leads to an overall negative climatic water balance (Fabig, 2007).

Land cover in Saxony-Anhalt is dominated by intensive agriculture (56%), with forests covering 23% (Statistisches Landesamt Sachsen-Anhalt, 2024). Most forests are even-aged monocultures of *Pinus sylvestris*, while native deciduous species, including *Quercus robur, Fagus sylvatica*, and *Alnus glutinosa* are restricted to fragmented remnants, often located in floodplain areas. Natural forest ecosystems currently represent only 16% of the forested area (Suck et al., 2014).

Saxony-Anhalt can be subdivided into distinct landscape units. The northern and northeastern lowlands, shaped by the Saale glaciation, feature sandy soils, pine heaths, and shallow groundwater tables, such as in the Ohre-Drömling and Dumme depressions. In contrast, the south and southwest transition into mid-mountain forelands is underlain by consolidated rocks (e.g., sandstone and limestone), with generally low groundwater productivity (LHW Sachsen-Anhalt, 2012). The central lowlands are dominated by the floodplains of the Elbe, Saale, and Mulde rivers, with complex alluvial and groundwater dynamics. Overall, 90% of the extracted groundwater comes from unconsolidated sandy aquifers in the north and east, likely supporting the occurrence of GDV (Jordan & Weder, 1995).

For this study, we focussed on (semi-) natural vegetation and thus excluded agricultural areas, artificial surfaces and water bodies using high-resolution regional land cover maps (MWL, 2023; Malinowski et al., 2020). Additionally, the Harz Mountains were excluded from the study due to a lack of data on groundwater depth. This might have excluded potential GDV such as mountain meadows or wetlands. The final study area is reduced by 60% and covers 8251 km^2^, representing vegetated landscapes across geological and climatic gradients.

### 2.2 Predictor variables

We selected eight predictor variables to represent vegetation physiology and vitality, as well as environmental controls on GDV distribution (e.g., Rampheri et al., 2023; Gomes Marques et al., 2019; Doody et al., 2017; Eamus et al., 2015). These variables were grouped into two categories: i) remote sensing predictors that refer to vegetation water use and greenness, and ii) geospatial predictors that control groundwater availability.

#### 2.2.1 Remote sensing predictors

Analyses were conducted for the meteorological summer (June–August), when temperate vegetation reaches maximum canopy development (Yang et al., 2017). In contrast to frameworks for arid and semi-arid regions that rely on distinct seasonal droughts, we assessed dry-year anomalies and interannual variation by comparing vegetation vitality between dry and wet summers as identified from long-term climatic anomalies.

First, to delineate dry and wet years, we used precipitation and temperature data (daily data at 1 km resolution) from the Germany National Meteorological Service (DWD, 2024a, b). Anomalies were calculated relative to the 1991–2020 climate baseline for each hydrological year (September–August). Meteorological summers with highly negative precipitation anomalies were defined as dry years (2018, 2019, 2020, 2022), while those with positive precipitation anomalies were defined as wet years (2017, 2021, 2023, 2024) (Figure 1).

**Figure 1.**
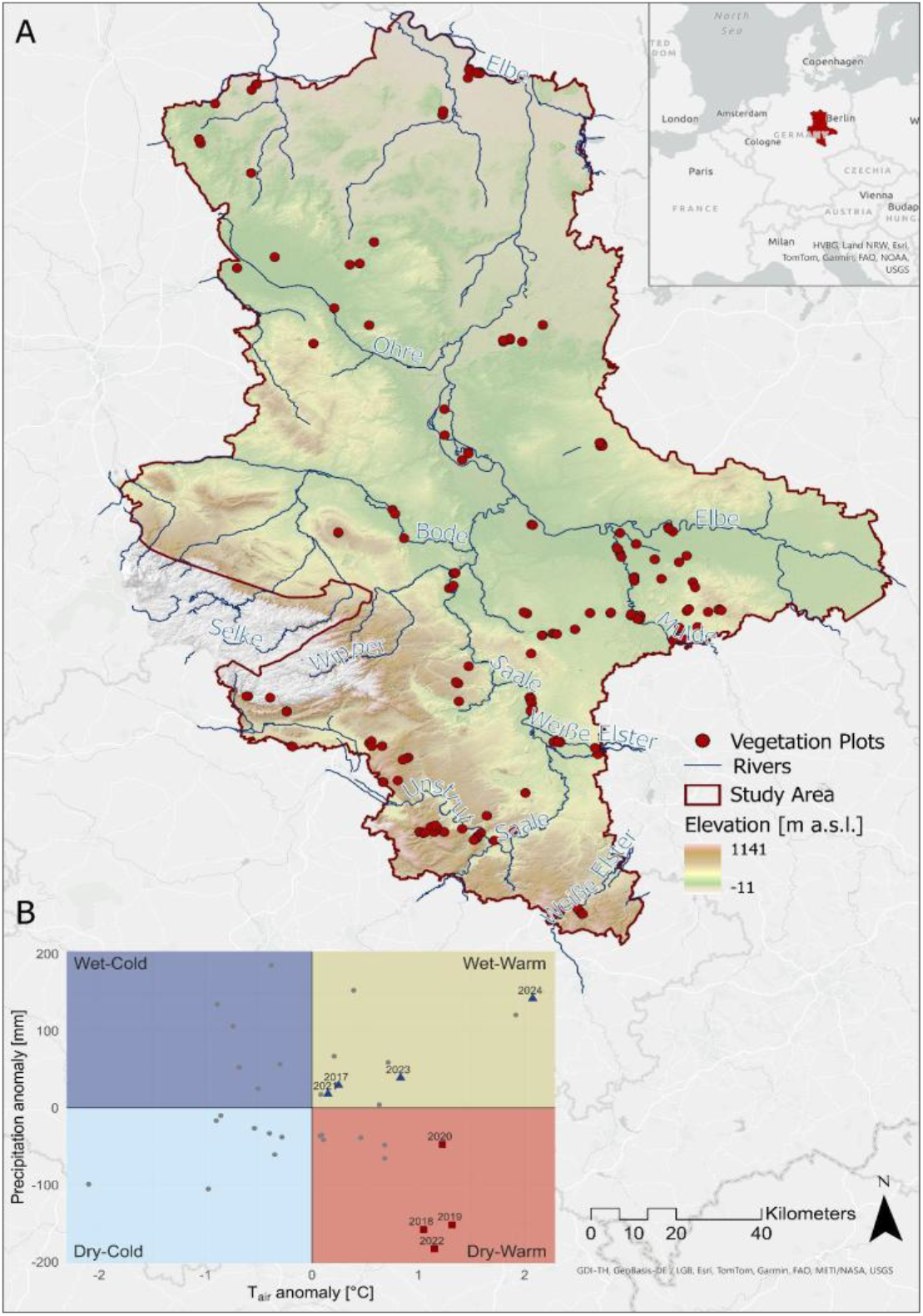
Overview map of the study area Saxony-Anhalt, Germany showing the elevational gradient and the locations of the 181 vegetation surveys. Panel B shows Precipitation (mm) and Temperature (°C) anomalies for Saxony-Anhalt calculated from DWD climate data for the reference period 1991-2020 (DWD, 2024a, b).

Second, we integrated ET data from ECOSTRESS (Hook & Fisher, 2019) together with vegetation indices derived from Sentinel-2 to detect GDV. ET provides a direct measure of water use by vegetation and is therefore a sensitive indicator of groundwater access, as GDV maintains high and stable ET even under drought stress (Gao et al., 2023; Chiloane et al., 2022; Eamus et al., 2016). ECOSTRESS retrieves land surface temperature and emissivity using a physics-based temperature and emissivity separation algorithm (TES) which forms the basis for generating higher-level products such as ET (Fisher et al., 2020). For this study, we used the Level-3 daily latent heat flux product at 70 m resolution (ECO3ETPTJPL), processed with the Priestley–Taylor Jet Propulsion Laboratory (PT-JPL) algorithm (Fisher et al., 2020; Priestley & Taylor, 1972). ET daily data was available for June–August 2018-2022, filtered for coverage (>30% of the study area), cloud-masked, and cleaned for extreme values (outside 8th–98th percentile) leaving 253 scenes for analysis (Supplementary 1).

From this dataset, we derived two ET-based predictors to identify ‘cool and wet islands’ of GDV (Barron et al., 2014): i) ETdry is the mean ET during the dry summers of 2018, 2019, 2020, and 2022. It was normalised by subtracting the mean ET per land cover class (Malinowski et al., 2020) and dividing it by the standard deviation per class to control for vegetation-type-specific ET variation (z-transformation); ii) ETdiff, the absolute difference in mean ET between the driest (2022) and wettest (2021) summers indicating interannual stability. These predictors provide insights into water use, its stability across hydrological and climatic variation, and the relative behaviour within vegetation types, providing a proxy of GDV occurrence.

Third, we quantified vegetation greenness using the Enhanced Vegetation Index (EVI) (Huete et al., 2002; Eq. 1). EVI was calculated for 786 Sentinel-2 scenes between 2017-2024 with a spatial resolution of 10 m in Google Earth Engine (Gorelick et al., 2017). Cloudy pixels were masked using the QA60 band.

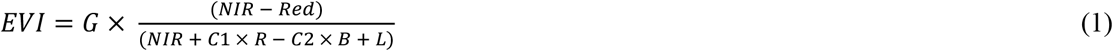

L is the canopy background adjustment. C1 and C2 are used to account for aerosol influences. Although the coefficients were originally developed for MODIS data, they are also widely used for Sentinel-2, as follows: L = 1, C1 = 6, C2 = 7.5, and G = 2.5 (Huete et al., 2002; Liu & Huete, 1995).

Finally, we calculated two vitality predictors to detect GDV following the ‘green island’ concept (Akasheh et al., 2008): i) EVIdry, the mean summer EVI across dry years, representing sustained greenness under drought; and ii) EVIsd, the interannual variability (standard deviation) in mean EVIdry between wet and dry years, where low values indicate consistent vitality and potential groundwater access. All pixels with EVI values < 0.2 were excluded from the analysis as they do not represent healthy vegetation (Huete et al., 2002).

#### 2.2.2 Geospatial predictors

We included three commonly used topographic predictors to capture environmental controls on groundwater availability and consequently, GDV distribution (e.g., Duran-Llacer et al., 2022; Gomes Marques et al., 2019; Doody et al., 2017). Elevation, slope, and the Topographic Wetness Index (TWI) were derived from a 5m digital elevation model (LVermGeo, 2023) and serve as proxies for topographically driven water accumulation (Münch & Conrad, 2007). The TWI was calculated in QGIS using Equation 2 (Beven & Kirkby, 1979):

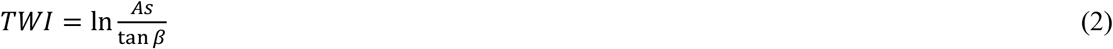

where *As* is the catchment area and *β* is the local slope angle.

We calculated groundwater table depth (GWTD) as key control on GDV distribution (Huang et al., 2019; Wierda et al., 1997) from regional groundwater contour maps (MWU, 2024). Groundwater contours were interpolated into a 10m raster surface, from which GWTD was calculated by subtracting the groundwater height from DEM-derived surface elevation. Where GWTD exceeded land surface elevation, it was adjusted to the surface value. Pixels with shallow GWTD within rooting depth are more likely to contain GDV due to easier access to groundwater (Ciruzzi & Loheide II, 2021).

### 2.4 Botanical field mapping

During the summers of 2023 and 2024, we recorded 181 vegetation plots (10 m × 10 m) across the study area, to generate ground-truth data for spatial modelling. Area-proportional random sampling aimed to reflect the areas of hydrogeological units (Thiele et al., 2023). Yet, aquitards were underrepresented due to the applied landcover masking. For each plot, species composition and relative cover were recorded using the Braun-Blanquet cover scale (Braun-Blanquet, 1928). Ellenberg-type Indicator Values (EIV) (Ellenberg, 1974) were assigned to each species using a European dataset (Tichý et al., 2023). Then, each recorded species was classified as obligate, facultative, or non-phreatophyte, based on a global phreatophyte and groundwater-associated species checklist (El-Hokayem et al., 2026). Since facultative phreatophytes can introduce uncertainty when used as indicators of GDV (Hultine et al., 2020), site-specific refinement was necessary. Following the approach of Lenkenhoff & Rose (2002), facultative phreatophytes occurring on sites with shallow groundwater table depth (GWTD < 5 m) were reclassified as site-specific phreatophytes, reflecting likely groundwater access. As groundwater tables below 5 m substantially reduces groundwater influence on tree growth and transpiration and hence indicates site specific non-phreatophytes (Ciruzzi & Loheide II, 2021). The ecohydrological classification for each plot was based on the site-specific phreatophyte cover (P%) as a proxy for GDV (El-Hokayem et al., 2026).

### 2.5 Classification of GDV

We classified GDV and non-GDV areas using Random Forest, an ensemble machine learning algorithm that builds multiple decision trees from random subsets of data (Breiman, 2001). Random Forest is widely used in remote sensing to map vegetation and groundwater-dependent ecosystems due to its robustness and capacity to model non-linear relationships (Rohde et al., 2024b; Rapinel et al., 2019; Pérez Hoyos et al., 2016).

As the pairwise Pearson correlation analysis was not significant, all eight predictor variables were included in the modelling (Supplementary 2). We applied both Random Forest classification and regression models. For binary classification, the ground-truth vegetation plots were classified as GDV or non-GDV based on predefined phreatophyte cover thresholds of 25%, 50%, and 75% representing statistical quartiles (El-Hokayem et al., 2026). For regression, phreatophyte cover was used as a continuous response variable. 15 ground-truth plots were excluded from the dataset due to missing values for ETdry. The final ground-truth dataset consisting of 166 plots was split into 80% training and 20% testing subsets. Then, hyperparameter tuning was conducted by adjusting the number of trees, the number of variables per split, minimum leaf size, bag fraction, and the maximum number of nodes hierarchically. This resulted in 108 model runs per phreatophyte cover threshold. Final model parameters were selected based on minimising Root Mean Square Error for regression and maximising overall accuracy for classification. The final Random Forest model was built using 50 trees, 2 variables per split, one minimum leaf population, 0.5 bag fraction and the maximum number of nodes available (Supplementary 2). The importance of the explanatory variables was extracted based on the Gini impurity index.

### 2.6 Plausibility analysis

We compared our mapped GDV areas to an independent biotope map (LAU, 2009). This map was derived from the interpretation of CIR aerial orthophotos from 2009, and categorises land cover into major units such as forest, woodland, herbaceous vegetation, water bodies, non-vegetated areas, agricultural, horticultural, viticultural, and built-up areas. These units are further subdivided into 42 structural units, each representing specific biotope or land use types with associated dominant plant species.

We classified twenty structural units as likely GDV based on either their inherent groundwater reliance (e.g., swamp forests, fens, tall forbs, springs) or dominance of phreatophyte species (*Alnus glutinosa, Salix* spp., *Populus* spp.). The classification, supported by literature (Kilroy et al., 2008; Wierda et al., 1997; Batelaan et al., 2003; Londo, 1975) and expert consensus of the authors, ensured an independent reference for plausibility analysis. We compared the classified GDV maps with these reference GDV biotopes to extract true positives and false negatives and calculated Producer’s Accuracy to quantify model performance.

## 3 Results

### 3.1 Vegetation plot analysis

The 181 vegetation surveys recorded 223 plant species, including 61 site-specific phreatophytes of which 24 and 37 were obligate and facultative. The highest proportion of phreatophytes occurred in the first tree layer (58%), and lower proportions in the second tree layer (24%), shrub layer (26%), and herb layer (17%). The most abundant phreatophyte species in the study area are: *Fraxinus excelsior, Quercus robur, Sambucus nigra, Alnus glutinosa*, and *Prunus padus*. Comparing EIV for GDV and non-GDV plots classified at the 25% phreatophyte cover threshold shows distinct ecological preferences (Figure 2). While GDV plots are more often associated with moist (median moisture value = 6.6) and nutrient rich soils (median nutrients value = 6.2), non-GDV plots indicate drier (median moisture value = 5.5) and less fertile soils (median nutrients value = 5.6). Also, while non-GDV plots are almost never found on saline soils, GDV plots can tolerate some salinity.

**Figure 2.**
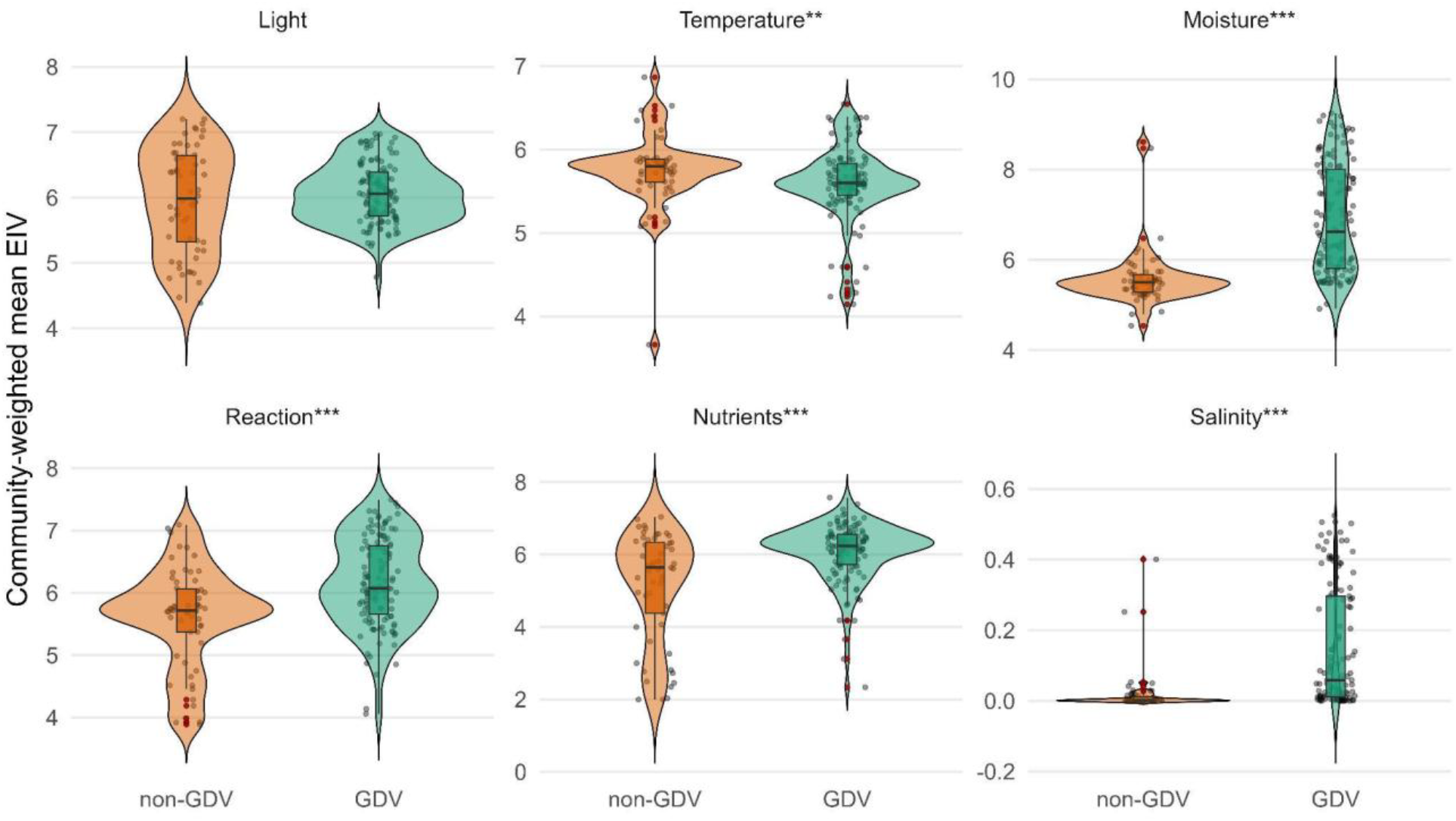
Community-weighted Ellenberg Indicator Values for 181 vegetation plots from the field surveys classified as non-GDV and GDV. Wilcoxon rank-sum tests show significant differences (asterisks) between GDV and non-GDV plots for all EIV except light availability. Red dots represent outliers in the data.

**Figure 3.**
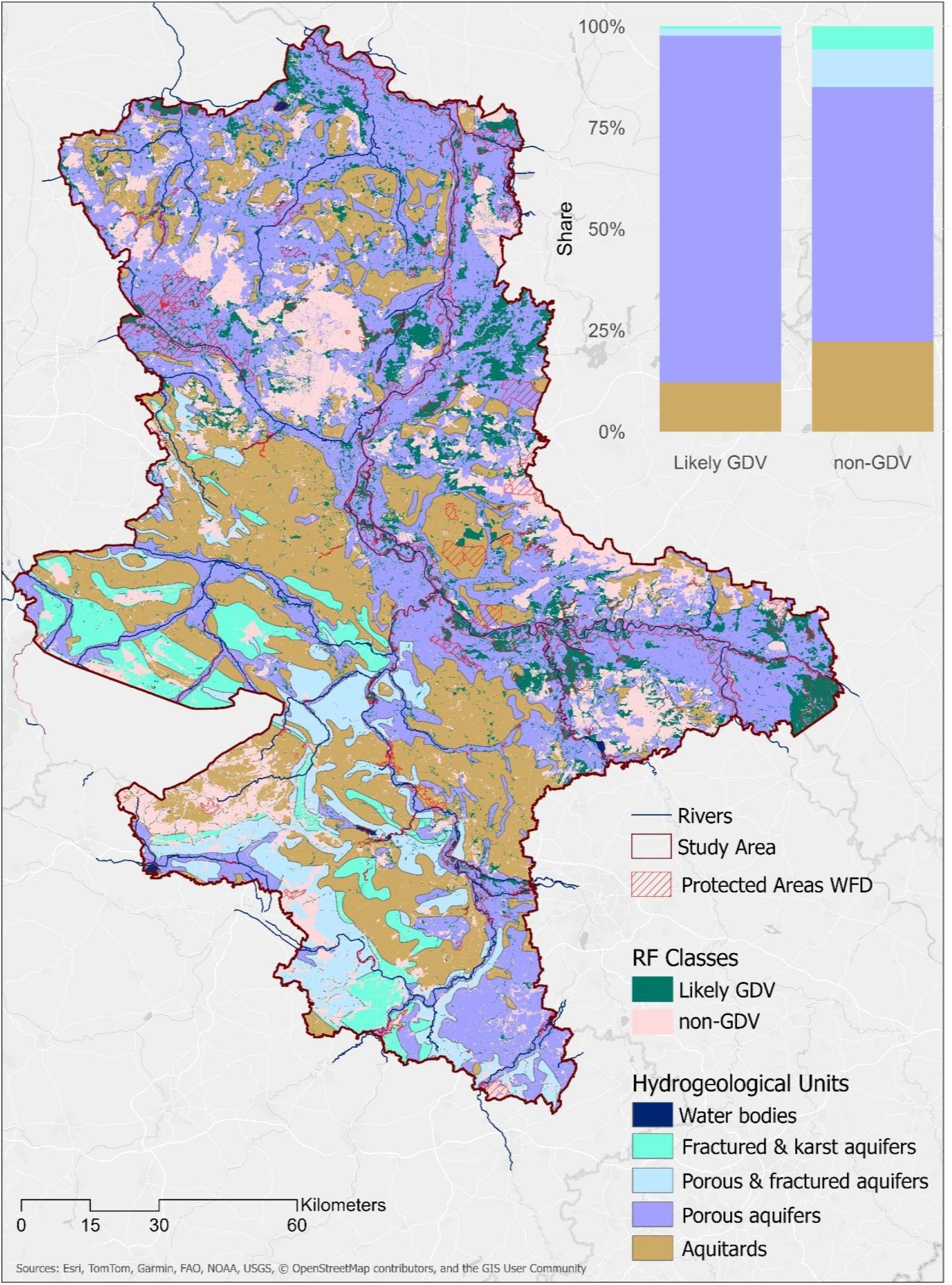
Results of the Random Forest (RF) classifier using the 25% phreatophyte cover threshold showing the distribution and extent of likely GDV and non-GDV in the study area and across distinct hydrogeological units. The map also displays protected areas according to the EU Water Framework Directive. Alternative maps using phreatophyte thresholds of 50% and 75% and Random Forest regression are shown in Supplementary 3.

### 3.2 GDV mapping and model evaluation

Approximately 2067 km^2^ of likely GDV areas (representing 41% of all analysed pixels) were mapped across Saxony-Anhalt at 10m resolution. From these classified GDV areas, 52% are located within protected areas, from which approximately one third are protected under the WFD. GDV occurrence is mainly associated with floodplain forests along the major rivers Elbe and Saale, and lowland depressions such as the “Ohre-Drömling”. Large contiguous areas of non-GDV occur in upland regions such as the Harz mountains and their forelands or heaths such as the “Dübener Heath” or the “Colbitz-Letzlinger Heath”. The Random Forest model using a phreatophyte cover threshold of 25% achieved the highest testing accuracy of 0.97. GWTD was the most important predictor, followed by elevation, TWI, and ETdiff. Plausibility analysis against a regional biotope map confirmed the reliability of predicted GDV areas, with 85% of GDV areas matching mapped alluvial, fen, and groundwater-fed biotopes.

The explanatory variables used in the Random Forest models highlight distinct environmental and ecological gradients across the study area (Supplementary 3 for detailed maps). Elevation peaks in the Harz Mountains, where steep slopes contrast sharply with the flatter lowlands of central and northern regions. This topographic gradient corresponds with hydrogeological patterns as TWI values are highest in the Elbe and Mulde river valleys and northern lowlands, indicating areas of potential water accumulation, while uplands show lower TWI values due to increased runoff. Similarly, groundwater table depth (GWTD) is generally shallow across the region, with approximately 44% of pixels showing GWTD < 5 m, decreasing with elevation.

Differences in ET between the summers of 2021 (dry) and 2022 (wet) are most pronounced around the city of Halle, the southern Harz Mountains, the “Dübener Heath” near Dessau, and the Middle Elbe floodplain. However, ETdiff shows relatively low spatial variability across the study area, with notably low differences in the “Dumme Depression” and north of the Havel River in northeastern Saxony-Anhalt. Land-cover-normalised ETdry exhibits a southeast-to-northwest gradient with highest ET values in the “Ohre-Drömling” and “Dumme Depression”, the “Altmark” lowlands, and floodplain forests along the northern Elbe and the Saale River. The spatial and temporal variability of EVI reflects drought and vitality patterns. EVIsd ranges from 0 to 0.35, with low variability overall, but higher interannual variability occurs in the southern Harz, mountain forelands, “Dübener Heath”, and fragmented northeastern forest patches. EVIdry exhibits spatial patterns closely mirroring those of ETdry, indicating areas of sustained vegetation vitality concentrated in river floodplains, low-lying depressions, and forested mountain foothills.

## 4 Discussion

### 4.1 Ecological patterns and management implications

Botanical field surveys revealed significant ecological differentiation between GDV and non-GDV communities. Temperate GDV favoured slightly cooler, wetter, nutrient-richer, and mildly acidic to alkaline environments compared to non-GDV. However, differences in species composition and EIV for moisture and nutrients were less pronounced than those observed between semi-arid GDV and non-GDV communities in the Mediterranean biome (El-Hokayem et al., 2023a). This likely reflects the more stable hydrological regime in temperate climates, where access to soil water remains high and there are no annual summer droughts (Xu & Wu, 2025). In semi-arid and arid environments, strong seasonality and pronounced water deficits promote the dominance of deep-rooted, obligate phreatophytes such as *Banksia littoralis, Quercus douglasii, Phoenix dactylifera*, or *Tamarix spp*. (Canham et al., 2012; Eamus et al., 2015; Chimner & Resh, 2014; Thomas, 2014) creating clearer ecological boundaries between GDV and non-GDV. In contrast, temperate GDV in the study area are typically composed of mixed stands of facultative phreatophytes, such as *Fraxinus excelsior, Alnus glutinosa*, and *Quercus robur*, whose groundwater use is opportunistic and varies with location (Orellana et al., 2012).

The mapped distribution of GDV in Saxony-Anhalt reflects regional hydrogeology and land use. GDV clusters predominately along river valleys, floodplains, and lowland depressions characterised by shallow groundwater and alluvial substrates, while upland areas with deep groundwater show sparse GDV occurrence. This pattern supports that GDV occurrence is primarily driven by the accessibility of the capillary fringe, with vegetation in low-relief and glaciated northern plains maintaining a hydraulic connection to groundwater during droughts (Doody et al., 2017). Conversely, elevated terrains and intensively cultivated loess plateaus exhibit few likely GDV areas due to both deep groundwater and agricultural drainage (Baaner et al., 2025). Similar spatial preference of GDV into floodplain and valley systems has been observed i.e. in semi-arid Australia and South Africa (Rampheri et al., 2023; Doody et al., 2017; Münch & Conrad, 2007). The overall proportion of GDV detected in Saxony-Anhalt with 41% of (semi-) natural vegetation is comparatively high for a temperate region, where 18-30% groundwater uptake are estimated across the entire biome (Link et al., 2023; Evaristo & McDonell, 2017). This might reflect the dry, continental climate in the study area, which may favour the development of GDV (Hultine et al., 2020), as well as strong hydrological connectivity in glacial and fluvial deposits especially in northern lowlands (Eismann et al., 2002). In the intensively used landscape of Saxony-Anhalt, only a small area of (semi-) natural vegetation remains in areas unsuitable for agriculture, such as depressions, wetlands and ravines. In turn, these landforms favour the occurrence of GDV, where they act as refugia for biodiversity and ecosystem functions under increasing climatic stress (Kløve et al., 2014; Münch & Conrad, 2007).

From a management perspective, our GDV map provides spatially explicit information relevant to the EU WFD, which seeks to protect groundwater-dependent ecosystems (Batelaan et al., 2003). The mapped GDV area is around 6,6 times larger than the area identified on the biotope map (313 km^2^). This suggests a significant amount of previously unmapped and potentially unprotected GDV. Anyhow, most mapped GDV in Saxony-Anhalt currently lie outside protected areas, indicating that these ecosystems are still underrepresented in conservation planning. Linking the GDV map to groundwater body assessments could prioritise ecologically vital resources, guide abstraction limits, and designate GDV sites, advancing policy integration in increasingly drought-prone temperate landscapes (Baaner et al., 2025; Pandey et al., 2023; Krogulec, 2018; Rohde et al., 2017).

The observed ecological and spatial patterns highlight the azonal character of GDV, determined by groundwater availability and accessibility rather than by climatic or edaphic zonation (Sieben, 2019). While the species composition and ecological traits of GDV communities differ between temperate and arid regions, their distributional patterns seem comparable overall. This suggests that fundamental ecohydrological controls on GDV occurrence (Glanville et al., 2023; Cantonati et al., 2020) might be consistent across biomes and thus comparably detectable through remote sensing and geospatial data integration (Pérez Hoyos et al., 2016). Yet, mapping frameworks must be adapted to regional climatic variability and data availability.

### 4.2 Methodological evaluation and outlook

Mapping temperate GDV is challenging due to the absence of distinct dry seasons, which complicates the detection of drought-related vegetation responses (Akasheh et al., 2008). To address this, we identified dry and wet years from precipitation anomalies, allowing interannual comparisons of vegetation water stress and ET dynamics as indicators of groundwater use. However, the transferability to wetter regions needs to be further tested. Projected increases in drought frequency and severity under climate change (Gebrechorkos et al., 2025; Kornhuber et al., 2019) may improve the applicability of such approaches in temperate regions, while at the same time increasing pressure on GDV (Kløve et al., 2014).

The use of satellite-derived ECOSTRESS ET data represents a novelty in GDV-mapping, providing a more direct indicator of vegetation water use. Our results show that high ET during dry conditions is a stronger predictor of GDV occurrence than interannual ET variability, suggesting that stable water use under drought is a useful signal of groundwater dependence (Sommer et al., 2016; Adams et al., 2015; Yang et al., 2011). This is consistent with findings for vegetation vitality from arid and semi-arid regions, where vegetation vitality during dry periods was more important in predicting GDV occurrence than interannual changes (El-Hokayem et al., 2026, Rohde et al., 2024b). In contrast to vegetation indices from satellite data, which often show lagged responses to climate extremes up to three months (Xu et al., 2026), ET responds more directly to changes in water availability and therefore has the potential not only to indicate, but ultimately to quantify, groundwater use by vegetation (Pierrat et al., 2025; Kulesza & Hościło, 2024; Macfarlane et al., 2017). However, forest management in the study region alters spectral signals, making groundwater detection difficult, particularly in managed pine and spruce forests. Also in such cases, ET-based indicators are advantageous as they are less sensitive to canopy disturbance. Still, high vegetation vitality does not always reflect groundwater use, since nutrient and light availability are important controls also (Brolsma & Bierkens, 2007).

GWTD was the most important predictor in our model, although its use in both field classification and spatial prediction requires acknowledging potential circularity. Regional patterns with shallow groundwater in northern lowlands and deeper GWTD in upland areas matched the spatial distribution of hydrogeological units and GDV, indicating that its predictive power extends beyond plot-level (Eismann, 2002). However, excluding GWTD reduced the testing accuracy from 0.97 to 0.78, demonstrating transferability to regions lacking detailed hydrogeological data while at the same time underscoring its ecological relevance. GWTD was also critical for ground-truthing temperate GDV, where widespread facultative phreatophytes complicate classification compared to arid regions where obligate phreatophytes dominate GDV (Hultine et al., 2019; Puertes et al., 2019; Evaristo & McDonell, 2017).

Interestingly, variable importance changed when increasing phreatophyte cover thresholds for the ground-truth plots, indicating a transition from hydrogeological controls to functional signals of groundwater use (Supplementary 2). At lower phreatophyte cover thresholds, GWTD dominated, reflecting likely GDV defined by groundwater accessibility, whereas with stricter thresholds ETdiff and ETdry became most important, suggesting that GDV with high phreatophyte coverage are characterised by differences in ET patterns than by groundwater availability alone. Overall, this pattern might suggest a transition from likely GDV (hydrogeological control) at low thresholds to functionally GDV (physiological control) at high thresholds. That said, conservative thresholds identify GDV by their active groundwater use rather than by environmental conditions alone, and underline the value of ET-based metrics for detecting GDV in temperate regions.

Despite methodological advances, several limitations remain for mapping GDV in temperate regions. Limited spatial coverage, coarse resolution (70 m), and short temporal record of ECOSTRESS restrict spatial transferability, fine-scale mapping and long-term analyses. Furthermore, cloud cover reduces the availability of optical Sentinel-2 and ECOSTRESS data, particularly in wet years, constraining interannual comparisons. Under warm, dry conditions, calculated ET values can exceed values in wetter years, which may cause mean ET to appear lower during cold and wet periods. Normalising ET to potential ET (i.e. using relative ET) would likely provide a more robust indicator of vegetation water use, although variability in satellite overpass timing would remain a challenge. The integration of radar data (e.g. Sentinel-1) could improve temporal coverage and provide more robust indicators of vegetation structure and plant water use in future studies (Castellazzi et al., 2024). A further limitation is the limited ground-truth dataset with 166 vegetation plots, which may constrain model generalisation. Future studies could test the framework with reduced input data, such as habitat or biotope maps as alternative training sources.

## 5. Conclusion

This study presents the first high-resolution remote sensing-based framework for mapping GDV in temperate regions, addressing a major geographical gap in GDV research. By adapting frameworks originally developed for semi-arid and arid regions, we demonstrate that the underlying principles of

GDV detection, linking vegetation vitality and water use to groundwater access during dry periods, are per se transferable to temperate environments when climatic variability is accounted for. Integrating ECOSTRESS ET data with Sentinel-2 vegetation indices, hydrogeological, and topographic predictors enabled the identification of GDV in the study area. Our Random Forest model classified likely groundwater-dependent vegetation (GDV) and non-GDV areas with an overall accuracy of 0.97. It revealed that approximately 41% of semi-natural vegetation in Saxony-Anhalt is likely to be groundwater-dependent, with 1,754 km^2^ that were not identified from existing biotope maps. However, roughly half of these are located within protected areas, stressing the underrepresentation of GDV in current conservation frameworks. Variable importance analysis further indicated a shift from hydrogeological controls at low phreatophyte cover thresholds to functional indicators of GDV at high thresholds, underscoring the value of ET for identifying functionally GDV. Together, these findings establish a transferable, data-driven approach for temperate GDV mapping and provide a spatial baseline to support biodiversity and water policy implementation in Saxony-Anhalt under the EU WFD.

## Acknowledgement

The research was funded by the Federal State of Saxony-Anhalt via the MLU|BioDivFund. LE and CC acknowledge the support of the German Centre for Integrative Biodiversity Research (iDiv) Halle-Jena-Leipzig.

## Data availability

Data and codes are openly available at DOI: 10.5281/zenodo.18977164

