## Supplementary Material for "Mapping groundwater-dependent vegetation in temperate climates on the example of Central Germany"

### bioRxiv

Léonard El-Hokayem, David Emanuel Schulz, Christopher Conrad, 2026

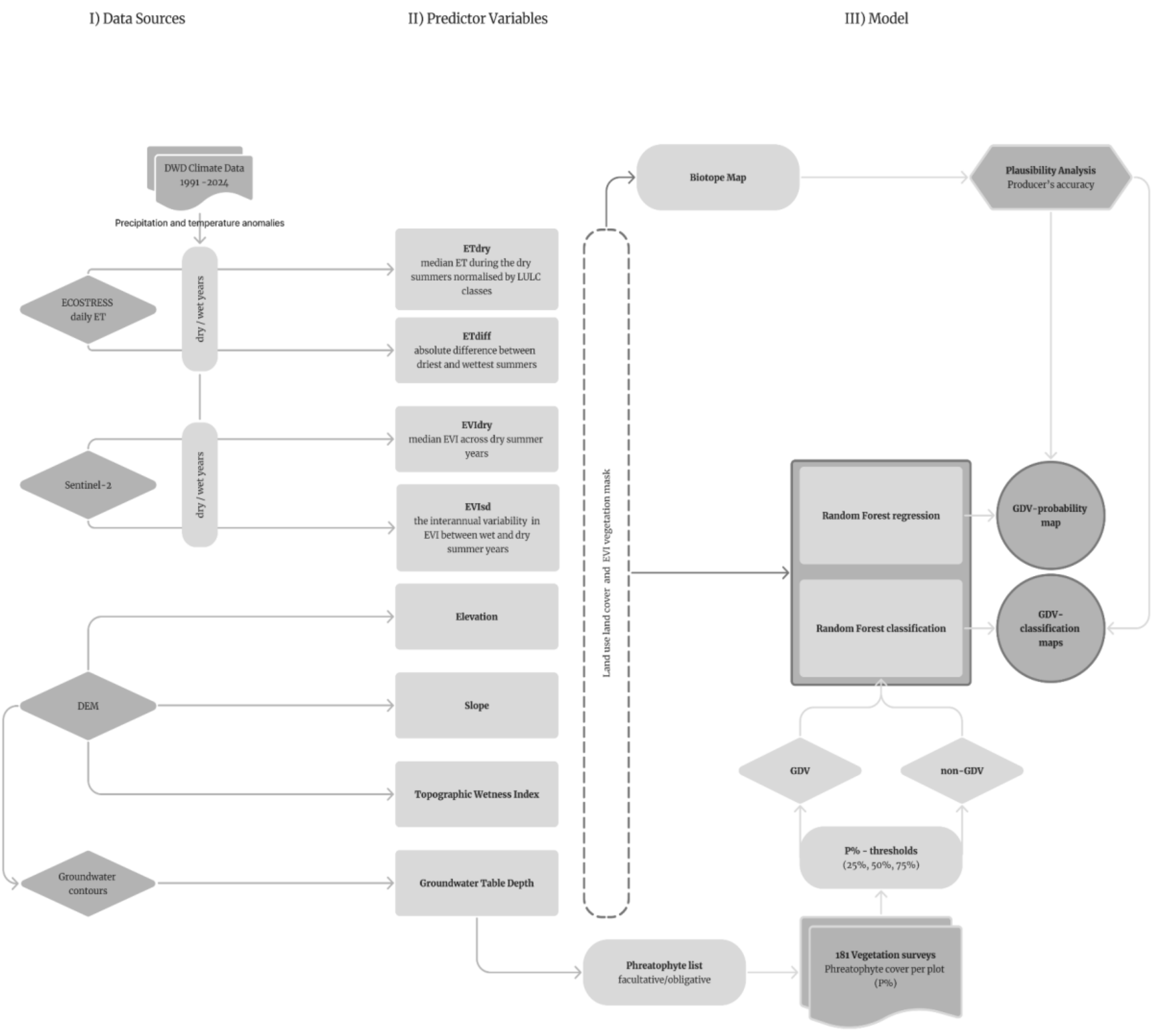


**Figure S1.1: Schematic overview of the mapping workflow.**

**Table S1.1: Overview of datasets used**

| Workflow | Variable | Dataset | Type | Spatial resolution | Temporal resolution | Period | Unit | Reference | Permanent Link |
|---|---|---|---|---|---|---|---|---|---|
| Temporal filter | Temperature | DWD | Raster | 1 km | daily | 1991-2024 | °C | DWD, 2024a | https://dwd-geoportal.de/products/GRD_DEU_P1M_T2M-N/ |
| Temporal filter | Precipitation | DWD | Raster | 1 km | daily | 1991-2024 | mm | DWD, 2024b | https://dwd-geoportal.de/products/GRD_DEU_P1M_RR/ |
| Mask | LULC | LULC Map Europe | Raster | 10 m | - | 2017 | - | Malinowski et al., 2020 | https://doi.org/10.3390/rs12213523 |
| Mask | EVIsummer | Sentinel-2 | Raster | 10 m | 5-days | 2017-2024 | - | Copernicus Sentinel data, 2024 | ee.ImageCollection("COPERNICUS/S2_SR_HARMONIZED") |
| Mask | Agricultural field margins | InVeKOS | Polygon | | annual | 2023 | - | MWL, 2023 | Not publicly available |
| Ground-truth | Vegetation surveys | | Point | 10 m | - | 2023-2024 | - | El-Hokayem & Schulz, 2026 | |
| Predictor | ET | ECOSTRESS LEVEL-3 EVAPOTRANSPIRATION, L3(ET_PT-JPL) | Raster | 70 m | daily | 2018-2022 | W/m² | Hook & Fisher, 2019 | https://ecostress.jpl.nasa.gov/data |
| Predictor | EVI | Sentinel-2 | Raster | 10 m | 5-days | 2017-2024 | - | Copernicus Sentinel data, 2024 | ee.ImageCollection("COPERNICUS/S2_SR_HARMONIZED") |

| Workflow | Variable | Dataset | Type | Spatial resolution | Temporal resolution | Period | Unit | Reference | Permanent Link |
|---|---|---|---|---|---|---|---|---|---|
| Predictor | Elevation | DEM | Raster | 5 m | - | 2023 | m | LVermGeo, 2023 | https://www.lvermgeo.sachsen-anhalt.de/de/gdp-dgm5.html |
| Predictor | Slope | DEM | Raster | 5 m | - | 2023 | degree | LVermGeo, 2023 | https://www.lvermgeo.sachsen-anhalt.de/de/gdp-dgm5.html |
| Predictor | GWTD | Groundwater contours | Polyline | 10 m | - | 2023 | m | MWU, 2024 | https://registry.gdi-de.org/id/de.st/9c426126-1973-4f53- 8c56-75eb8e3f670e |
| Plausibility analysis | Biotopes | CIR 2009 Biotope Map | Polygon | 1:10,000 | - | 2009 | - | LAU, 2009 | Not publicly available |

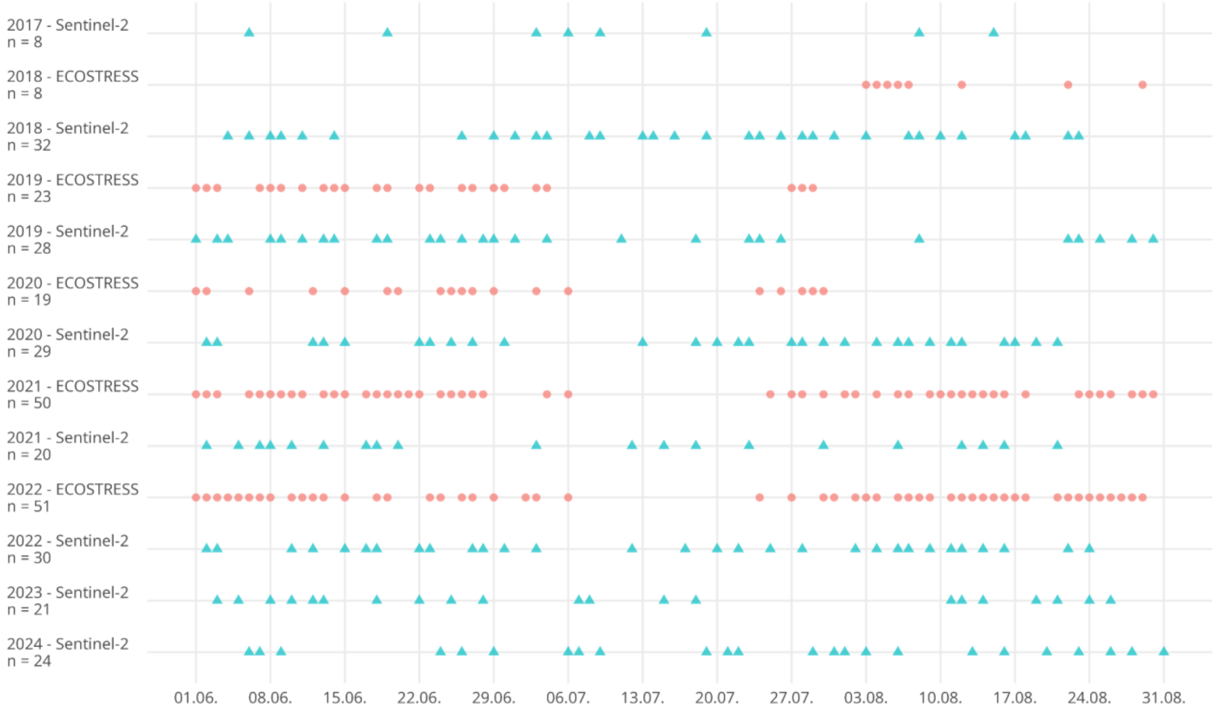


**Figure S1.2: Acquisition dates and number of available scenes for Sentinel-2 and ECOSTRESS data in the study area for the period 2017-2024.**

### Supplementary 2 Model details

**Mapping groundwater-dependent vegetation in temperate climates on the example of Central Germany**

**bioRxiv**

Léonard El-Hokayem, David Emanuel Schulz, Christopher Conrad, 2026

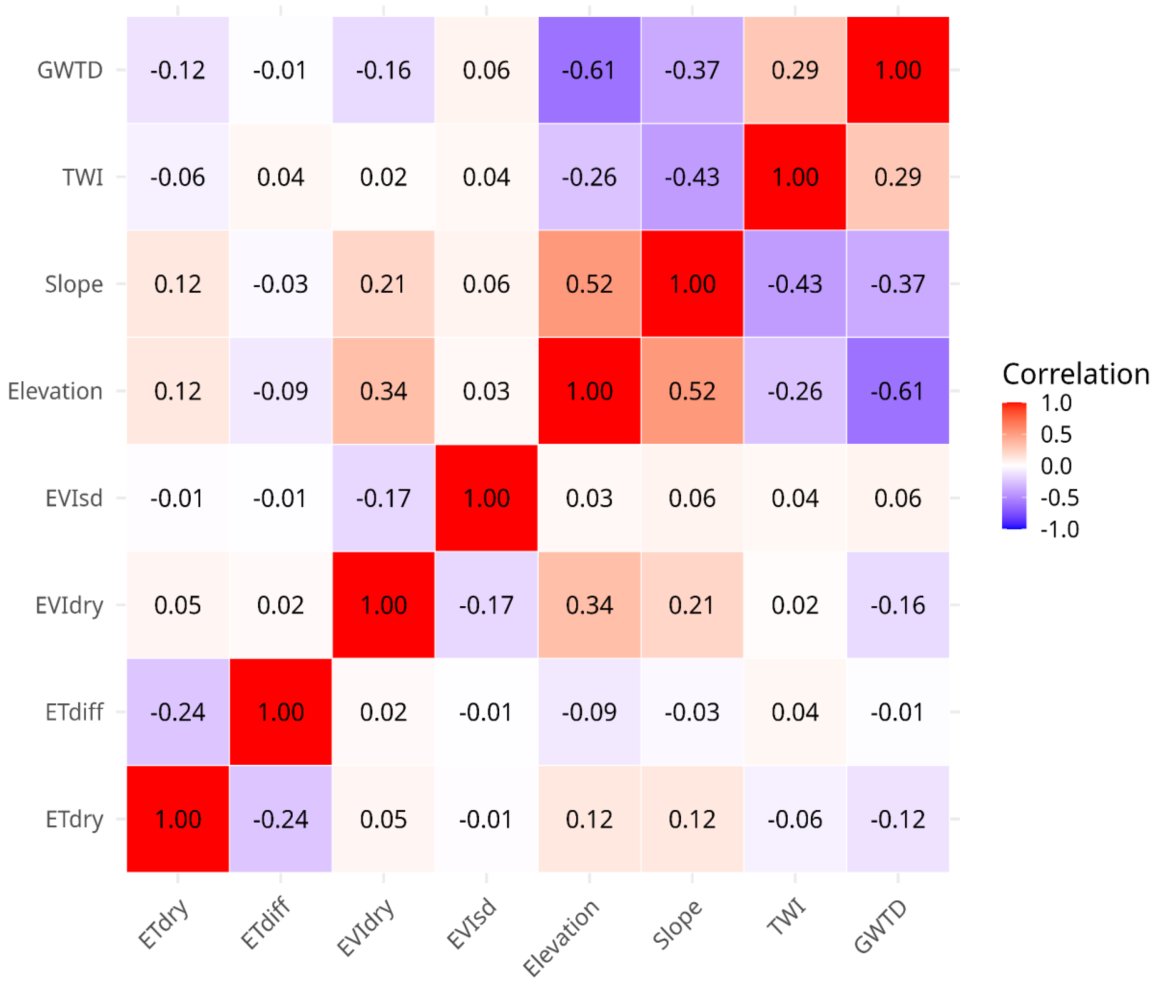


**Figure S2.1: Correlation matrix for eight indicator values extracted across the study area.**

**Table S2.1: Results of the hyperparameter tuning for the three Random Forest classifiers and the Random Forest regression model.**

| Model | N of trees | Variables per split | Min Leaf | Bag Fraction | Max Nodes | RMSE | Testing Accuracy |
|---|---|---|---|---|---|---|---|
| RF Classisification (P% 25) | 50 | 2 | 1 | 0.5 | max | - | 0.97 |
| RF Classisification (P% 50) | 20 | 2 | 2 | 0.5 | max | - | 0.89 |
| RF Classisification (P% 75) | 10 | 2 | 1 | 0.5 | max | - | 0.93 |
| RF Regression | 130 | 6 | 1 | 0.7 | max | 17.4 | - |

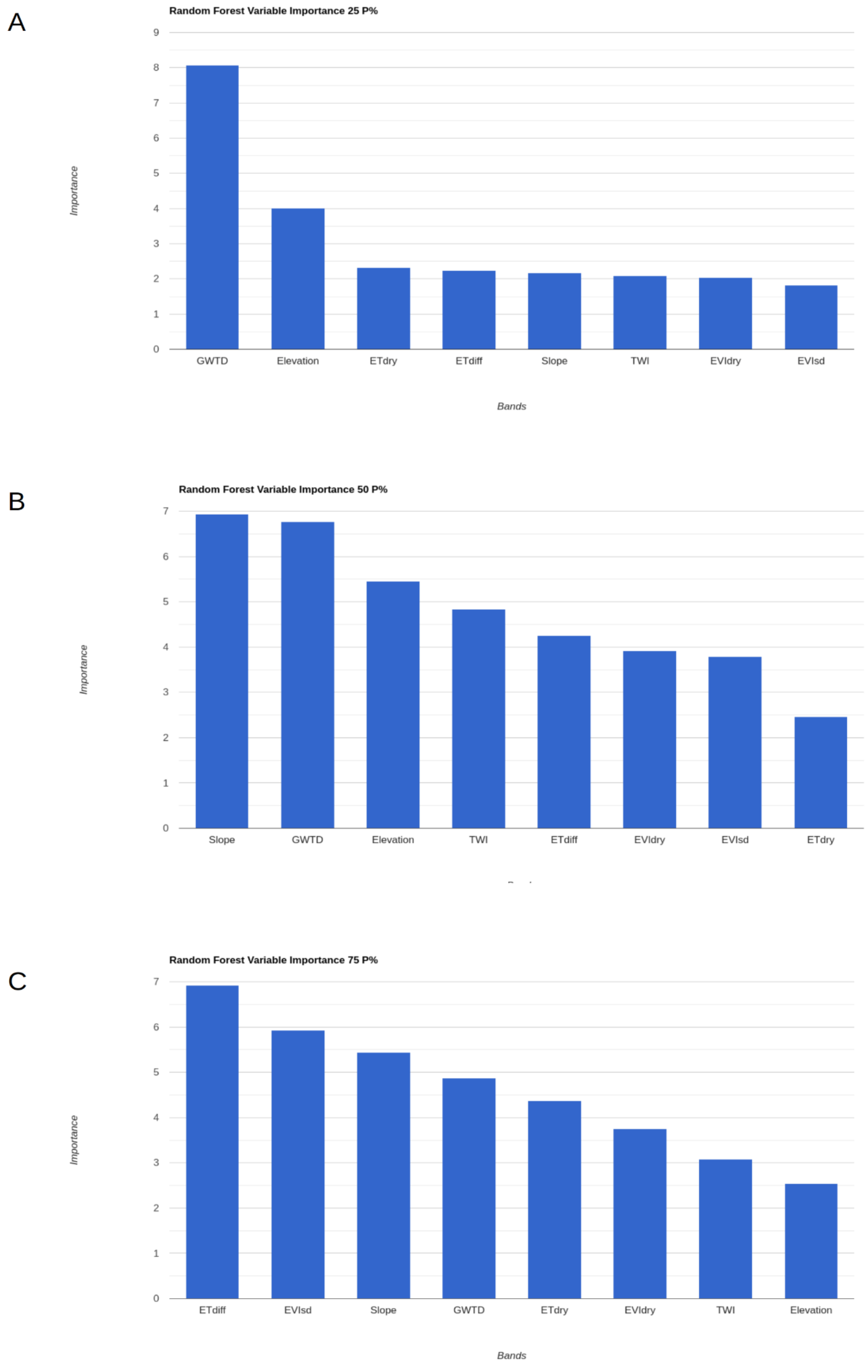


**Figure S2.2: Extracted variable importance of the Random Forest classifiers in Google Earth Engine with phreatophyte cover thresholds of A) 25%; B) 50%; and C) 75%. The importance is based on the Gini impurity index.**

### Supplementary 3 Result details

**Mapping groundwater-dependent vegetation in temperate climates on the example of Central Germany**

**bioRxiv**

Léonard El-Hokayem, David Emanuel Schulz, Christopher Conrad, 2026

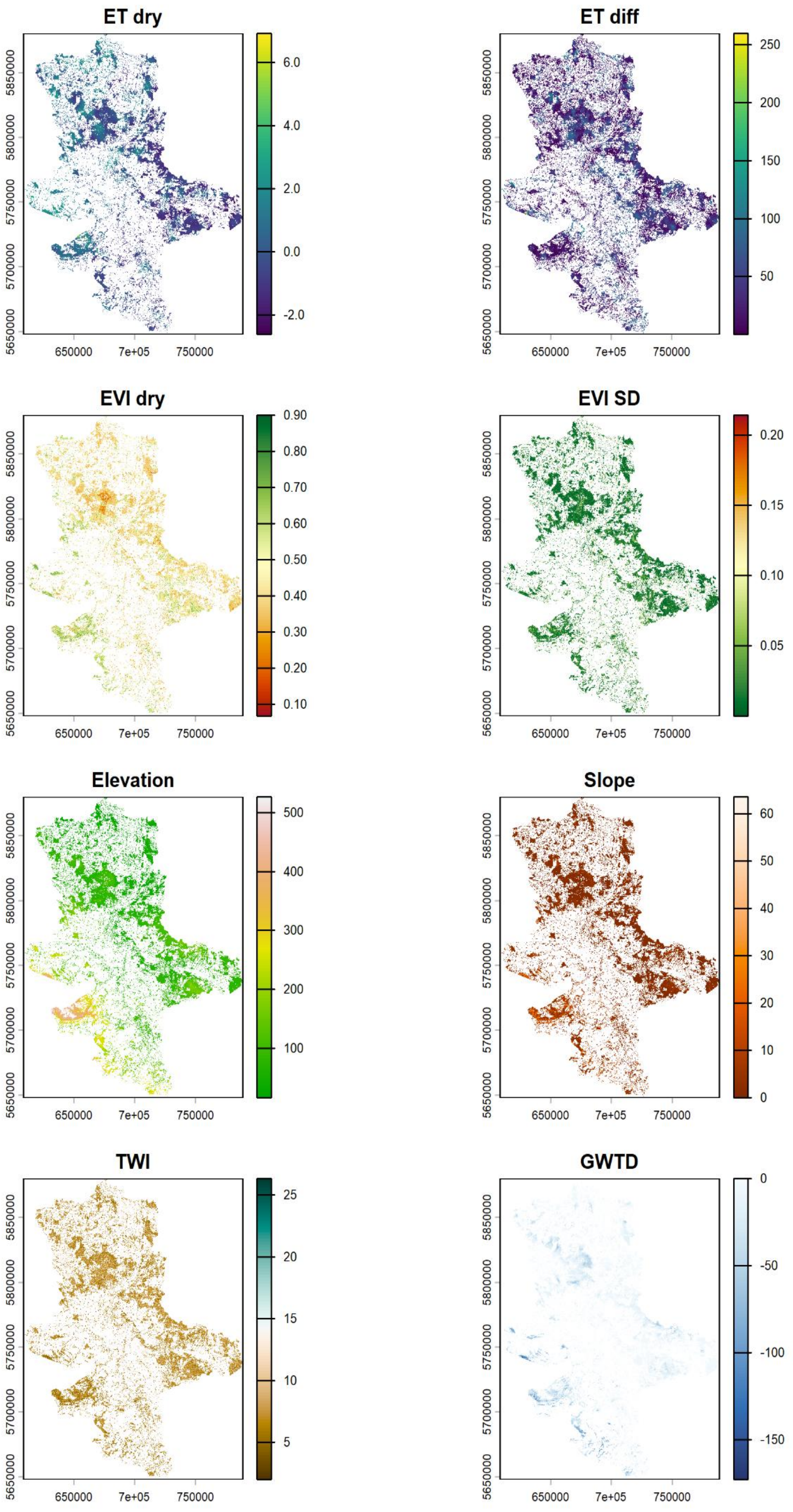


**Figure S3.1: Remote sensing and geospatial data products that were used as explanatory variables for the Random Forest models. With ETdry, ETdiff [W/m$^2$], EVIdry, EVIsd, elevation [m a.s.l.], slope [deg], TWI, and GWTD [m].**

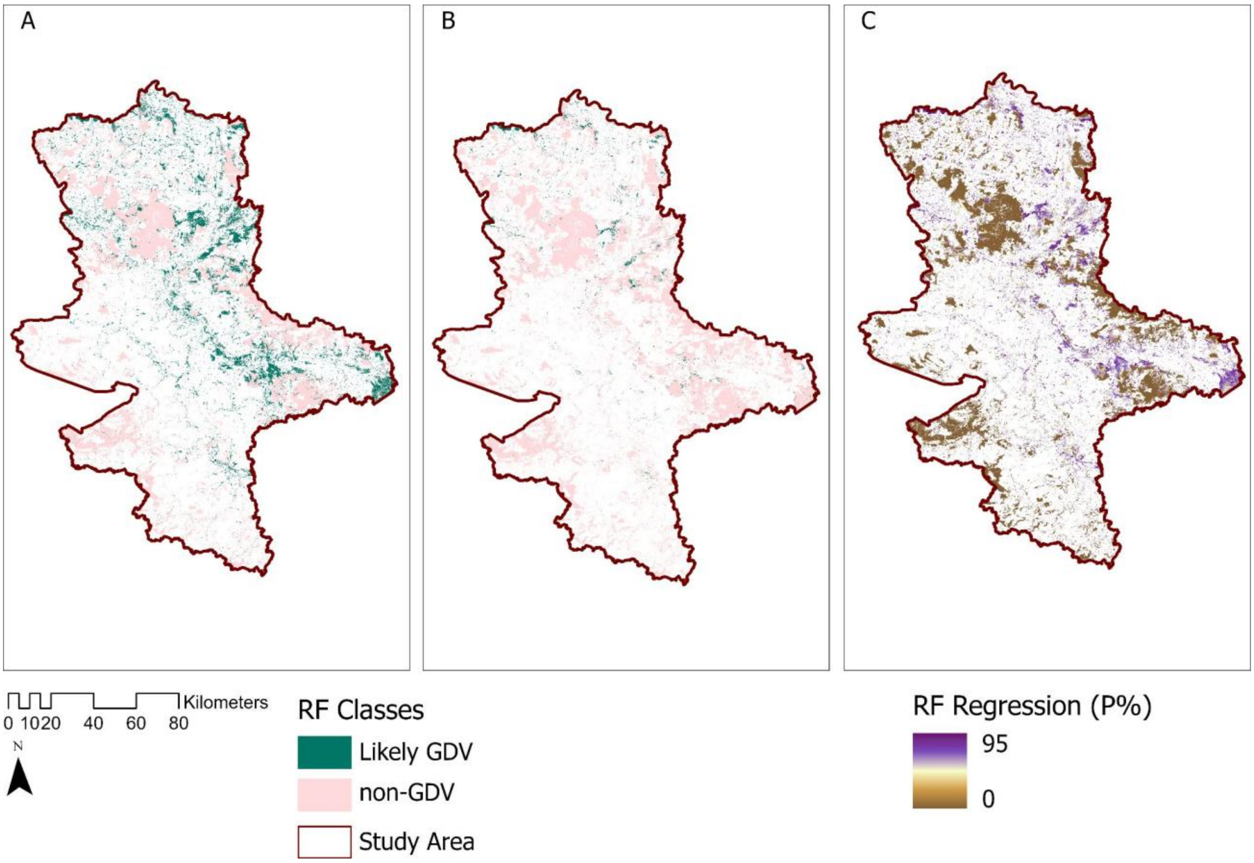


**Figure S3.2: Alternative GDV maps of the study area using phreatophyte cover thresholds of A) 50% with km² of likely GDV and B) 75% with km² of likely GDV. C) Result of the Random Forest regression using phreatophyte cover as continuous response variable.**
